# Binaural (pre)processing for contralateral sound field attenuation and improved speech-in-noise recognition

**DOI:** 10.1101/2021.01.22.427757

**Authors:** Enrique A. Lopez-Poveda, Almudena Eustaquio-Martín, Fernando Martín San Victoriano

## Abstract

Understanding speech presented in competition with other sound sources can be challenging. Here, we reason that this task can be facilitated by improving the signal-to-noise ratio (SNR) in either of the two ears and that in free-field listening scenarios, this can be achieved by attenuating contralateral sounds. We present a binaural (pre)processing algorithm that improves the SNR in the ear ipsilateral to the target sound source by linear subtraction of the weighted contralateral stimulus. Although the weight is regarded as a free parameter, we justify setting it equal to the ratio of ipsilateral to contralateral head-related transfer functions averaged over an appropriate azimuth range. The algorithm is implemented in the frequency domain and evaluated technically and experimentally for normal-hearing listeners in simulated free-field conditions. Results show that (1) it can substantially improve the SNR (up to 20 dB) and the short-term intelligibility metric in the ear ipsilateral to the target source, particularly for speech-like maskers; (2) it can improve speech reception thresholds for sentences in competition with speech-shaped noise by up to 8.5 dB in bilateral listening and 10.0 dB in unilateral listening; (3) it hardly affects sound-source localization; and (4) the improvements, and the algorithm’s directivity pattern depend on the weights. The algorithm accounts qualitatively for binaural unmasking for speech in competition with multiple maskers and for multiple target-masker spatial arrangements, an unexpected property that can inspire binaural intelligibility models.

## INTRODUCTION

In daily listening scenarios, listeners often wish to attend to a sound source (the ‘target’) and ignore other, competing sound sources (the ‘maskers’). For example, while dining in a restaurant, a listener may wish to understand what a friend is saying disregarding other talkers. It is not yet totally clear how listeners with normal hearing can solve this so-called ‘cocktail party’ problem (Cherry, 1953), but it is well documented that hearing-impaired people have more trouble at this task than do people with normal hearing. The signal-to-noise ratio (SNR) required to achieve the same amount of speech understanding (e.g., 50% sentence recognition) is up to 9 dB higher for hearing-impaired than for normal-hearing people (e.g., Festen and Plomp, 1990; Peters et al., 1998). For this reason, assistive listening devices, including hearing aids and cochlear implants, often incorporate pre-processing algorithms that improve the SNR before the stimulus is further processed and delivered to the user. Here, we present a binaural (pre)processing algorithm that enhances the SNR in the ear ipsilateral to the target source by attenuating the contralateral sound field.

There exist many different pre-processing approaches to improve the SNR (for a recent review, see Doclo et al., 2015). Most algorithms assume that some knowledge is available *a priory* to distinguish the target from the masker sources. Some algorithms are based on assumptions about the temporal or spectral characteristics of the target or the maskers and enhance the SNR by filtering out the maskers in the time-frequency domain (e.g., Boll, 1979). Other algorithms, known as beamformers, are based on assumptions about the spatial location of the target or the masker sources and enhance the SNR by spatial filtering out the masker sources (e.g., Cox et al., 1987). (Beamformers typically involve using two microphones in each device to attenuate masker sounds from the rear.) Other algorithms can blindly separate the multiple sound sources in the acoustic scene and improve the SNR by attenuating the masker sources. Once separated, the target source needs to be selected by the user or based on assumptions about its location or acoustic characteristics. Blind-source separation requires as many microphones as sound sources are to be separated, including the user’s own voice (Hamacher et al., 2005; Doclo et al., 2015).

The approaches just described are monaural. For users of bilateral devices, monaural SNR-enhancement algorithms are often applied separately in each device, an approach that can distort inter-aural time (ITD) and level differences (ILDs) and thus hinder the localization of sound sources (e.g., Cornelis et al., 2012; Doclo et al., 2015). The introduction of assistive listening devices that swap signals between the two ears has spawned the development of binaural pre-processing algorithms, i.e., systems that couple the functioning of two devices (Hamacher et al., 2005; Moore, 2007; Baumgärtel et al., 2015a, 2015b). One approach consists of detecting binaural cues and re-introducing them after the SNR is enhanced separately in each ear (see the review of Doclo et al., 2015). Another approach involves taking advantage of the greater number of microphones available in the two devices to perform blind separation of a greater number of sources (binaural source separation with only one microphone per ear does not improve intelligibility when there are three masker sources; Luts et al., 2010). Liu et al. (2001) proposed an algorithm that could localize multiple sound sources in a scene with only two microphones and improve the SNR by cancelling the unwanted sources. As with monaural systems, however, a limitation of binaural source separation or localization algorithms is that the target still needs to be selected by the user or differentiated from the maskers based on assumptions (Hamacher et al., 2005; Doclo et al., 2015).

Users of assistive listening devices often have reduced sensitivity to binaural cues or no access to them. Indeed, (1) some hearing-impaired users have reduced sensitivity to ITDs for narrow-band stimuli (see Best and Swaminathan, 2019); (2) bilateral cochlear implants provide minimal access to ITDs (e.g., Grantham et al., 2008); and (3) the use of bilateral hearing aids and cochlear implants reduces or distorts ILDs when the two devices apply amplitude (or dynamic range) compression independently to each ear (e.g., Wiggins and Seeber, 2013; Lopez-Poveda et al., 2016a, 2019). Because binaural cues are important in ‘cocktail party’ listening situations (Bronkhorst and Plomp, 1989; but see Baltzell et al., 2020), some binaural pre-processing algorithms are designed to enhance naturally occurring binaural cues that may be reduced or absent for the listener. For example, some algorithms detect head-shadow ITDs, which are present at low frequencies (< 1500 Hz) and convert them into ILDs. These algorithms can improve sound source localization for users of bilateral hearing aids (Moore et al., 2016) and cochlear implants (Brown, 2018). They can also help bilateral cochlear-implant users recognize speech in competition with a speech interferer (Brown, 2014). However, they have failed to improve speech-in-noise recognition for bilateral hearing-aid users (Moore et al., 2016). One important limitation of this approach is that ILD enhancement per se needs not improve the SNR. To improve the SNR, ILDs should be enhanced separately for each sound source. Kokkinakis and Pirhosseinloo (2017) proposed an algorithm that enhanced the SNR for a target in competition with a single masker by enhancing the ILD for the masker alone. This algorithm, however, requires knowing the specific location of the masker as well as the listener’s head-related transfer function (HRTF) for the masker location, and this information needs not be available a priori.

Here, we propose a binaural (pre)processing algorithm that improves the SNR in the ear ipsilateral to the target by attenuating the contralateral sound field. The algorithm is inspired by the binaural cochlear-implant sound processing strategy of Lopez-Poveda et al. (2016a, 2016b, 2017, 2018, 2020). That strategy involved dynamic contralateral control of the compression in each frequency channel so that the stronger spectro-temporal features in each ear can suppress themselves in the contralateral ear. As a result, a sound source is enhanced in the ipsilateral ear but attenuated in the contralateral ear. In cocktail-party scenarios with multiple talkers, the target talker is unlikely centered (Grange and Culling, 2016) and listeners can switch attention to the ear that has the better SNR for the momentary target (Bronkhorst and Plomp, 1988; Brungart and Iyer, 2012). Therefore, in cocktail party listening scenarios, the strategy of Lopez-Poveda et al. (2016a) can improve speech intelligibility without making any assumptions about the location of the target source^1^.

The algorithm proposed here was intended to produce a similar effect as the strategy of Lopez-Poveda et al. by subtraction of the weighted contralateral stimulus. The weight is regarded as a free parameter, but we propose setting it equal to the ratio of ipsilateral to contralateral HRTFs for an anthropometric manikin averaged over an appropriate azimuth range (see below). The algorithm is implemented in the frequency domain and evaluated technically and experimentally for listeners with normal hearing.

## THE ALGORITHM

In natural listening situations, our brain has access only to the pressure waveforms sensed by the left and right eardrums. In a free-field acoustic scenario where a target sound source, *S*(*t*), is presented in competition with *n* masker sources [*M_i_*(*t*), for *i*=1…*n*], the waveforms at the left and right ear, *L*(*t*) and *R*(*t*), respectively, can be expressed as a linear summation of the pressure waveforms generated by all sources, each convolved with the corresponding head-related impulse response (HRIR), as follows:

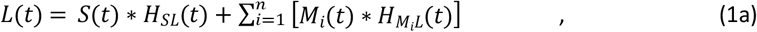

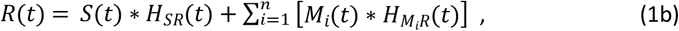

where *t* denotes time; *H_SL_*(*t*) and *H_SR_*(*t*) denote the HRIRs for the signal source for the left and right ears, respectively; *H_M_i_L_*(*t*) and *H_M_i_R_*(*t*) denote the HRIRs for the *i*-th masker, *M_i_*(*t*), for the left and right ears, respectively; and asterisks (*) denote the convolution operator.

Thanks to the properties of the Fourier transform, Eq. (1) may be expressed in the frequency domain as follows:

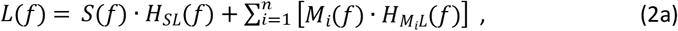

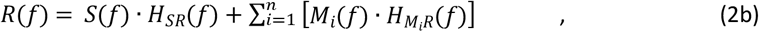

where *f* denotes frequency, each time waveform in Eq. (1) has been replaced by its corresponding complex frequency spectrum in Eq. (2), the HRIRs have been replaced with the corresponding frequency-domain complex HRTFs, and the convolution operator has been replaced by a product.

Let *L*′(*f*) and *R*′(*f*) denote the frequency spectra of the processed stimuli. The proposed approach consists of subtracting the weighted, unprocessed contralateral stimulus from the stimulus in each ear, as follows:

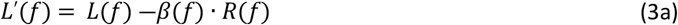

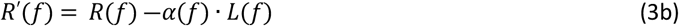

where *α*(*f*) and *β*(*f*) are the subtraction weights.

The subtraction weights are regarded as free parameters and complex numbers (i.e., with magnitude and phase). Therefore, they may be set to any appropriate values that improve the SNR. In the Appendix, it is mathematically shown that the values of *α*(*f*) and *β*(*f*) that would optimally cancel all maskers in the left and right ears would be as follows:

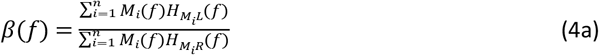

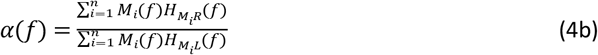

Equations (4a) and (4b) show that aiming for a perfect cancellation of the maskers at the two ears would be impractical because it would require knowing the spectrum of every masker, *M_i_*(*f*), as well as the HRTFs for each individual listener and for each masker location for the left and the right ear [*H_M_i_L_*(*f*) and *H_M_i_R_*(*f*)]. This information needs not be available a priori. Furthermore, in many listening situations, the identity and location of the signal and masker sources, thus the spectra of the maskers, change from moment to moment. For instance, in ‘cocktail party’ listening situations, the target and masker speakers often change during the conversation, and so do their locations relative to the listener as the listener moves the head and/or the target and masker talkers move around the listener. This makes it impractical, if not impossible, to calculate *α*(*f*) and *β*(*f*) for optimal cancellation of the masker(s) in an arbitrary listening situation.

Because *α*(*f*) and *β*(*f*) are free parameters, however, they can be set to *any* appropriate value for a specific purpose. In what follows, we set:

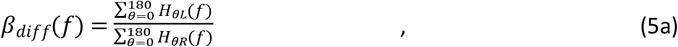

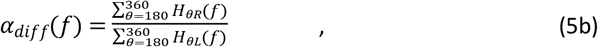

where *θ* denotes azimuthal angle, going from 0° for a source right in front to 360° clockwise, and *H_θL_*(*f*) and *H_θR_*(*f*) denote the complex HRTFs for an average listener. The appropriateness of this choice is discussed later. For convenience, in what follows, we will use the notation *β_diff_* and *α_diff_* rather than *β_diff_*(*f*) and *α_diff_*(*f*).

Note that for a symmetrical head:

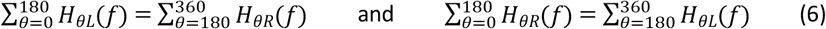

Hence:

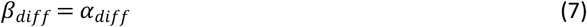

The rest of this paper is devoted to showing that this algorithm (Eq. 3) with *α_diff_* and *β_diff_* can improve the SNR in the ear contralateral to the masker source and thus intelligibility of speech in noise without significantly altering the lateralization of sound sources. Before doing so, however, several comments are in order.

1. We propose that *α_diff_* and *β_diff_* be calculated using average, rather than individual, HRTFs. In other words, our proposed approach does not require using individualized HRTFs. In what follows *α_diff_* and *β_diff_* were calculated for a KEMAR, an anthropometric acoustic manikin with average body dimensions (Burkhard and Sachs, 1975). Their magnitude and phase are shown in **Fig. 1**. The effect of calculating *α_diff_* and *β_diff_* using individualized and non-individualized HRTFs is analyzed and discussed below.
2. The proposed algorithm (Eq. 3) is linear. Therefore, the waveform that results from applying the algorithm to the signal+masker stimulus is equal to adding the waveforms that result from applying the algorithm to the signal and the masker separately, i.e.:

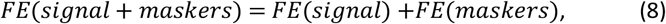

where *FE*(*x*) denotes the front-end algorithm applied to signal *x*. The linearity property is important because it implies that the algorithm works equally at all signal and masker levels, i.e., across SNRs.
3. *β_diff_* is the theoretical optimal *β*(*f*) to provide an infinite SNR (i.e., to perfectly cancel all maskers) in the left ear in a hypothetical listening scenario with infinite, identical [*M_i_*(*f*) = *M*(*f*) for all *i*] point-source maskers located at azimuthal angles from 0° to 180°. In other words, *β_diff_* is the optimal weight for the algorithm to provide an infinite SNR in the left ear for a ‘diffuse-field’ masker in the right hemifield. Similarly, *α_diff_* would be the optimal weight to deliver an infinite SNR in the right ear for a ‘diffuse-field’ masker in the left hemifield. With *α_diff_* and *β_diff_*, the algorithm does not provide perfect cancellation in any other condition, but still improves the SNR in many listening situations, as shown below.
4. The algorithm results in the directivity patterns shown in **Fig. 2**. (Note that the pattern is shown only for the left ear because the pattern for the right ear would be symmetrical about the midline.) The algorithm attenuates contralateral sound sources at low frequencies (0-1500 Hz), particularly at azimuths about 45° and 135°, and it slightly enhances contralateral high frequencies (>1500 Hz) at the same azimuths (**Fig. 2B**). This can enhance the SNR in the ear contralateral to the masker, for masker frequencies below about 1500 Hz.
5. The directivity pattern is similar for white noise (**Fig. 2A, B**) and speech-shaped noise (**Fig. 2C, D**). However, the overall SNR enhancement is expected to be greater for speech-shaped noise than for white noise because speech-shaped noise has more low-frequency content and less high-frequency content than white noise (compare **Fig. 3** and **Fig. 4**). The SNR enhancement effect of the algorithm for different masker types is analyzed and discussed below.
6. The algorithm attenuates contralateral low-frequency sounds with minimal changes in the level of ipsilateral sounds (**Fig. 2**). This distorts the head-shadow ILDs. This distortion and its effect on sound localization is analyzed and discussed below.

**Figure 1.**
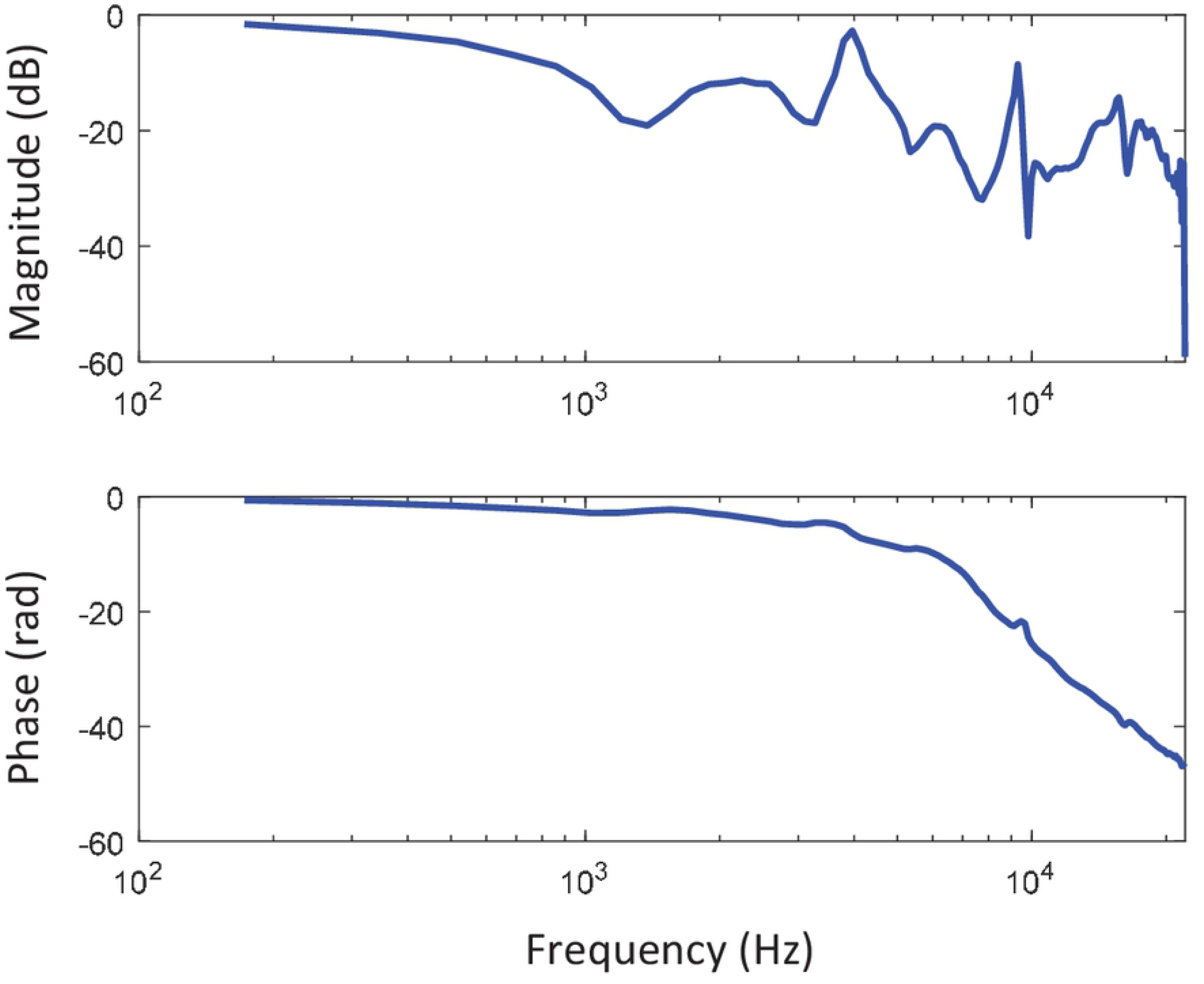
The magnitude (top) and phase (bottom) spectra of *α_diff_* and *β_diff_* (Eq. 5) calculated using the HRTFs for a KEMAR (Gardner and Martin, 1995). Note that *α_diff_* = *β_diff_* (Eq. 7) and that the phase is unwrapped.

**Figure 2.**
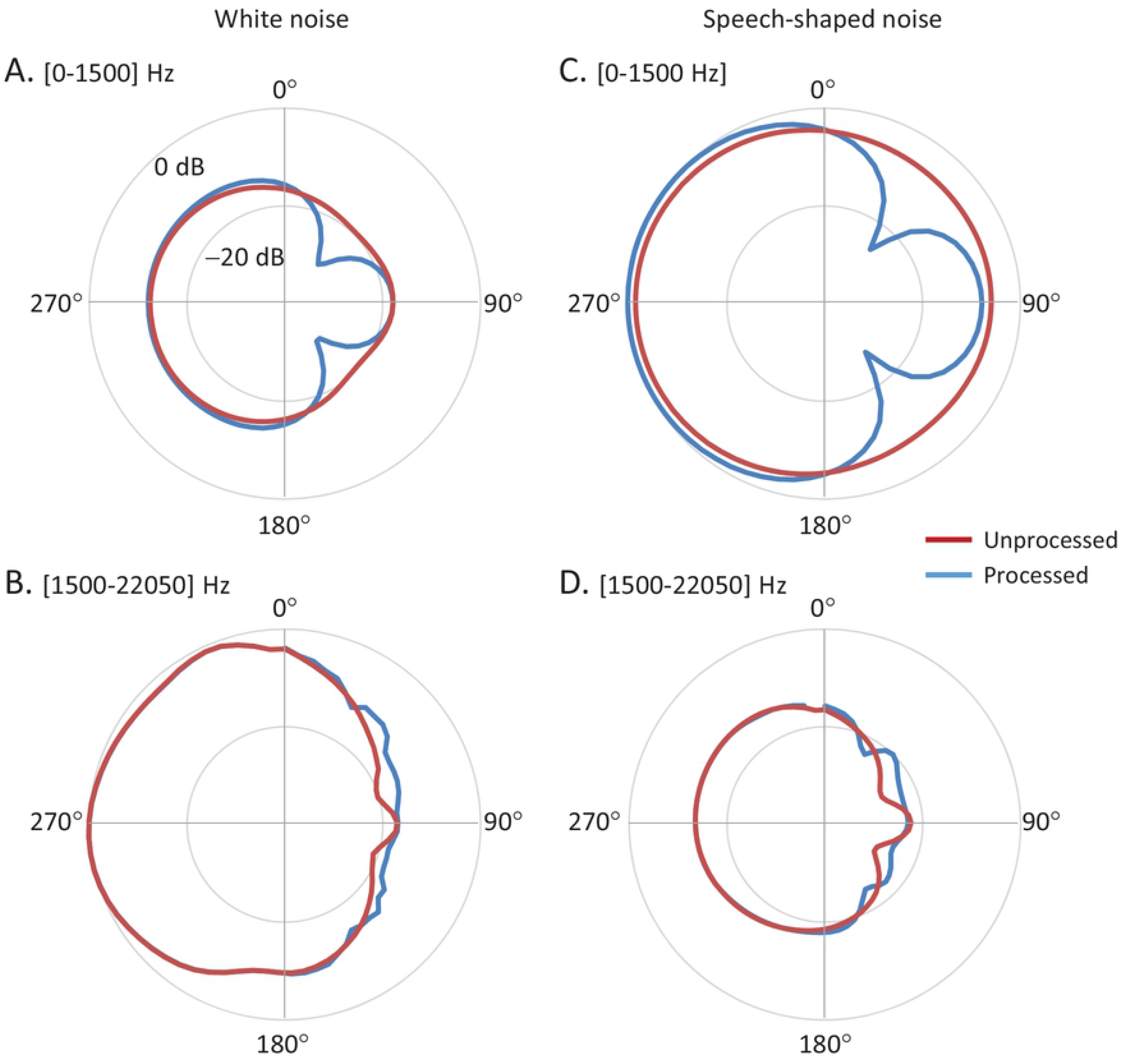
Directivity patterns for the left ear with and without processing using *β_diff_*. Patterns are shown for white noise (left panels) and speech-shaped noise (right panels). The top and bottom panels show patterns for the low- (0-1500 Hz) and high-frequency (>1500 Hz) content of each noise stimulus, respectively. The directivity pattern for the right ear would be symmetrical about the midline.

**Figure 3.**
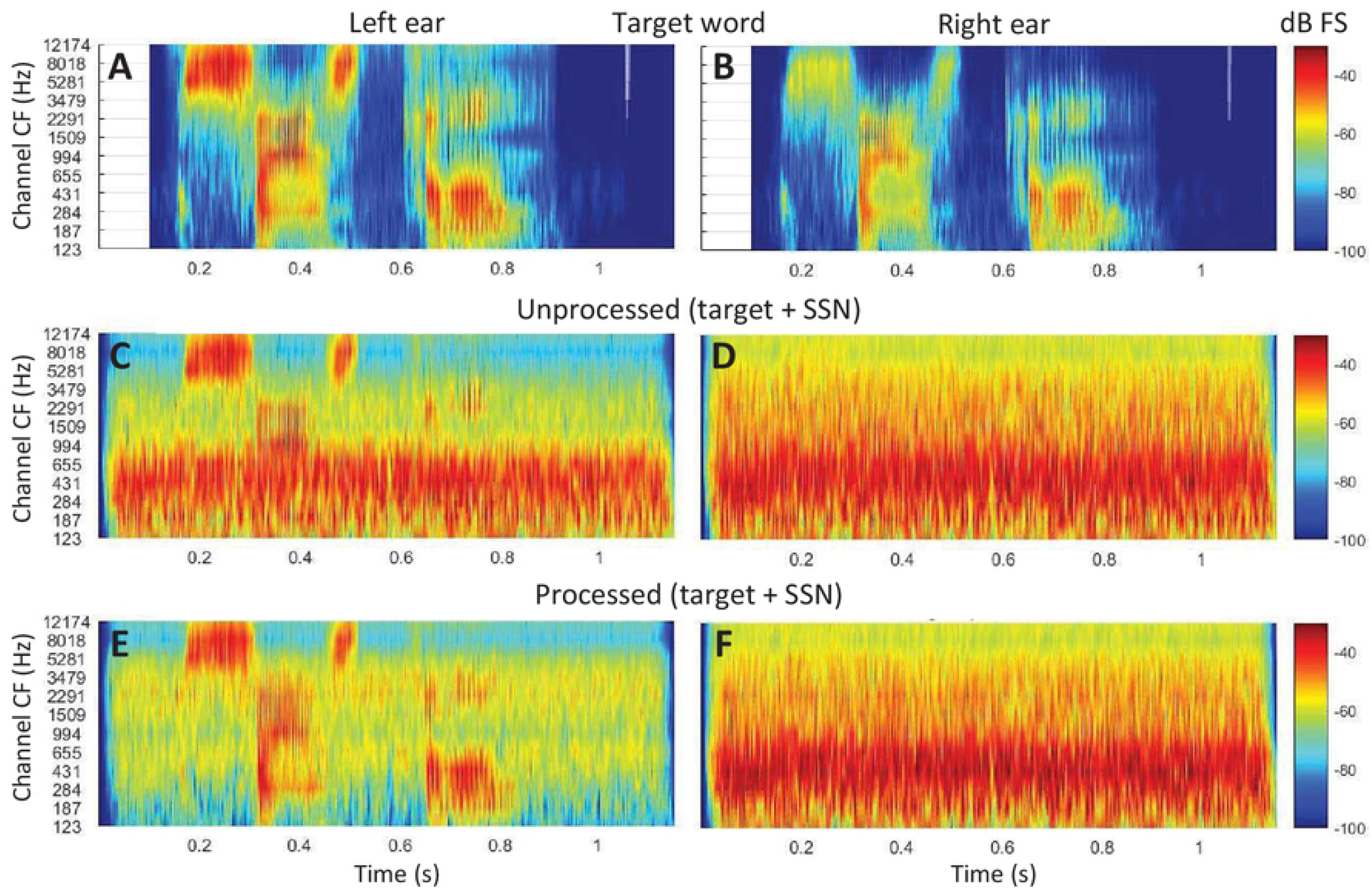
Example spectrograms for unprocessed (middle row) and processed (bottom row) stimuli in the left and right ears, respectively. The target was the Spanish word “sastre” at 270° azimuth in competition with a single source of speech-shaped noise at 45°. The SNR before HRTF filtering was −10 dB SNR. For reference, the top panels illustrate the spectrograms for the HRTF-filtered speech alone.

**Figure 4.**
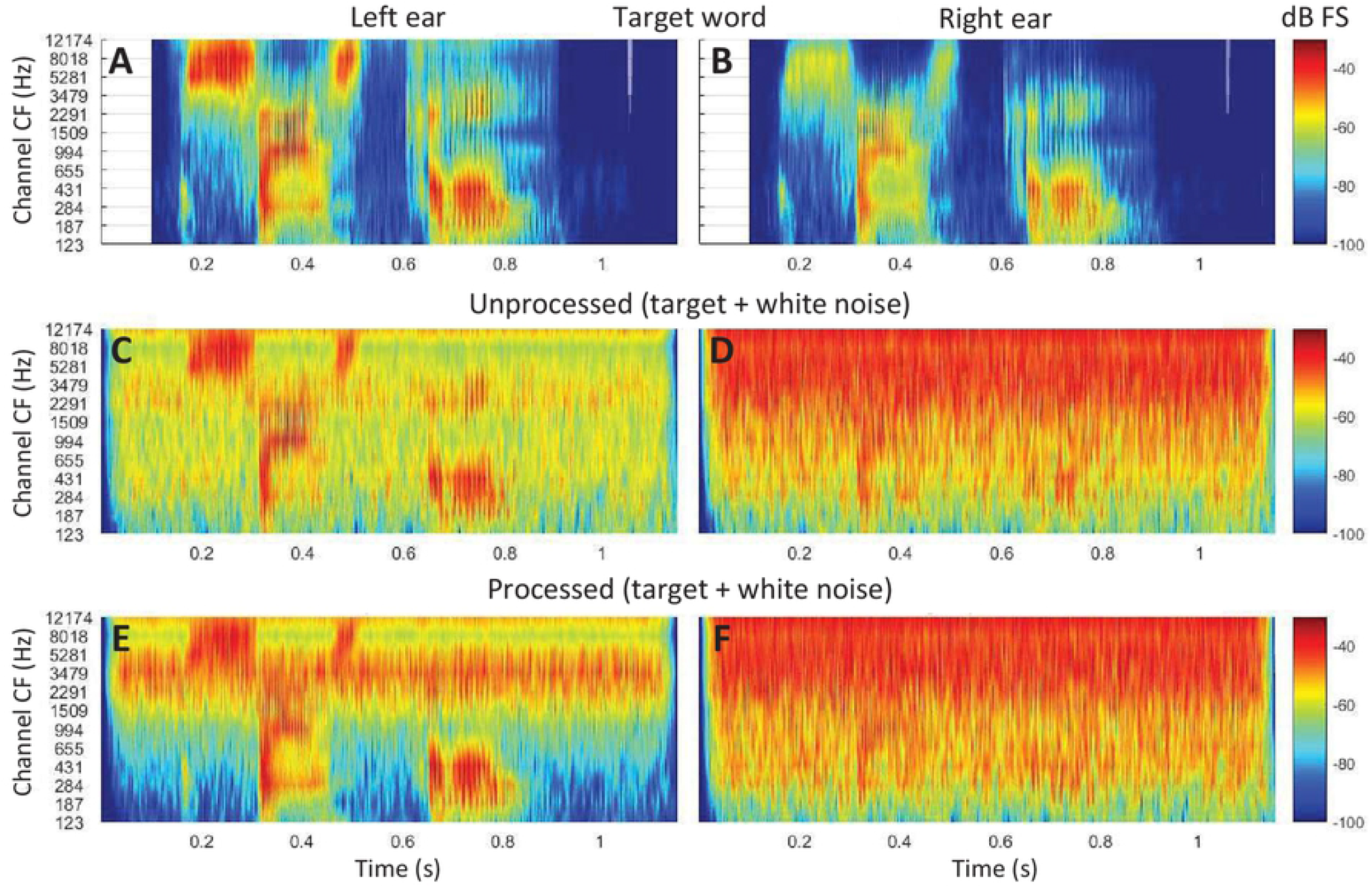
As **Fig. 3**, but for white noise rather speech-shaped noise.

## MATERIALS AND METHODS

### Evaluations

The proposed algorithm (Eq. 3) with *α_diff_* and *β_diff_* (Eq. 5) was evaluated technically and experimentally. The technical evaluation involved comparing the short-term objective intelligibility (STOI) (Taal et al., 2011) as well as the SNR for processed versus unprocessed stimuli. The experimental evaluation involved comparing the SNR at 50% speech recognition (termed speech reception threshold or SRT in noise) as well as sound-source lateralization performance in quiet for listeners with normal hearing for processed versus unprocessed stimuli. *α_diff_* and *β_diff_* (shown in **Fig. 1**) were always calculated using the HRTF database for a KEMAR (Gardner and Martin, 1995).

All evaluations were performed in a simulated free-field. All sound sources were at eye level (0° elevation). To achieve the desired target-masker spatial configurations, monophonic sound recordings were filtered through diffuse-field HRTFs. Unless otherwise stated, the same KEMAR HRTF database that was used to calculate *α_diff_* and *β_diff_* was also used to simulate free-field listening. This would be equivalent to calculating *α_diff_* and *β_diff_* using individualized HRTFs. For a subset of conditions (see below), however, we also assessed the benefits provided by algorithm when using non-individualized *α_diff_* and *β_diff_*. In this case, the spatial location of the sound sources was simulated by filtering stimuli through a different set of HRTFs (from IRCAM subject 1017^2^).

Experimental tests were approved by the Ethics Committee of the University of Salamanca. Participants were volunteers and not paid for their services. All of them signed an informed consent to participate in the study.

### Implementation

The algorithm was implemented in the frequency domain. Monophonic time-domain input waveforms for the target and masker(s) were first linearly scaled to the desired level. Here, levels are expressed in decibels relative to a maximum amplitude of unity and denoted as dB FS (full scale). The scaled, monophonic waveforms were convolved through appropriate HRIRs to obtain the unprocessed waveforms at the left and right ears, *L*(*t*) and *R*(*t*), for the desired (simulated) free-field target-masker spatial configuration. The stimulus waveforms were then subject to a frame-based processing to obtain their corresponding complex frequency spectra. To do it, the input waveform in each ear was divided into time-overlapping frames of 128 samples (the frame overlapping was 64 samples). Each signal frame was multiplied with a Hanning window and the result was padded with zeros to obtain a frame of 256 samples (64 samples of zeros were added before and after the signal frame). A fast Fourier transform (FFT) was applied to the windowed, zero-padded frame to obtain the complex frequency spectrum of the frame. The processing algorithm (Eq. 3) was applied to the resulting complex spectrum for each frame, and an inverse FFT was applied to obtain the processed waveforms, *L*′(*t*) and *R*′(*t*).

### Signal-to-noise ratio

For unprocessed stimuli, the SNR (in decibels) was calculated separately in each ear as follows:

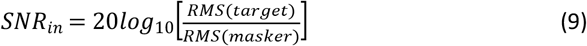

Where *RMS*(*target*) and *RMS*(*masker*) denote the root-mean-square (RMS) amplitude of the waveform in the ear in question for the HRTF-filtered target alone and the HRTF-filtered masker(s) alone, respectively.

Similarly, the SNR for processed stimuli was calculated in each ear as follows:

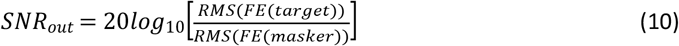

Where *FE*(*target*) and *FE*(*masker*) denote the waveforms at the output of the algorithm for the HRTF-filtered target alone and the HRTF-filtered masker(s) alone, respectively.

The SNR improvement (dB) provided by the algorithm was calculated as:

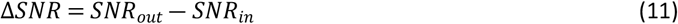

Positive ΔSNR values indicate that the algorithm improved the SNR (i.e., that the SNR was greater for processed than unprocessed stimuli) while negative values indicate the algorithm decreased the SNR.

Note that *SNR_out_* may be calculated using Eq. (10) because the algorithm is linear (Eq. 8). Also note that in the above calculations, the RMS amplitudes were calculated based on the input and output time waveforms to and from the processor, even though the algorithm was implemented in the frequency domain.

### Short-term objective intelligibility

The STOI is an objective measure of the amount of speech information available when the speech is degraded by a masker and/or processed by a sound processor (Taal et al., 2011). It is the average linear correlation over time and frequency between the unprocessed speech in quiet and the processed speech in noise. It is a scalar value between 0 and 1 that is expected to have a monotonic relation with the percentage of correctly understood speech tokens averaged across a group of listeners. Therefore, higher STOI values indicate higher SNR and better intelligibility in noise.

Here, the proposed algorithm was evaluated by calculating how much it improved the STOI.

The STOI improvement, ΔSTOI, was calculated as follows:

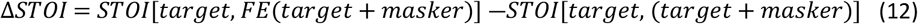

Where the notation *STOI*(*x*_ref_, *x*) indicates that STOI was calculated for stimulus *x* having stimulus *x*_ref_ as the reference. In other words, positive ΔSTOI values indicate that intelligibility, as predicted by STOI, would be greater for processed, *FE*(*target*+*masker*), than for unprocessed (*target*+*masker*) stimulus. STOI was calculated separately for the left and right ears having HRTF-filtered stimuli as the reference.

Unless otherwise stated, in calculating STOI, the target consisted of 10 concatenated sentences (with pre- and post-silence periods removed) from practice list #1 of the Castilian-Spanish hearing-in-noise test (HINT) (Huarte et al., 2008). STOI was calculated for three types of maskers: Gaussian white noise (WN), Gaussian noise filtered to have the long-term spectrum of speech (referred to as speech-shaped noise or SSN), and an international female fluctuating masker (IFFM) (Holube et al., 2010). The levels of the target and the masker were set equal to −20 dB FS (i.e., the SNR was 0 dB). The masker waveform was identical for processed and unprocessed stimuli to allow a fair comparison.

### Speech recognition tests

For listeners with normal hearing (NH), SRTs for sentences in noise were compared for processed and unprocessed stimuli. Speech reception thresholds were measured in bilateral and unilateral listening modes. Unilateral listening tests involved processing stimuli through the algorithm but stimulating only the ear ipsilateral to the target source (i.e., the left ear in this case, see below). Unilateral listening tests were aimed at testing to what extent ΔSNR (which is calculated for each ear) is a reasonable predictor of the actual SRT improvement.

#### Procedures

SRTs were measured using an adaptive procedure. The target was a sentence from the Spanish matrix sentence test (Hochmuth et al., 2012) and the masker was steady-state noise with the average long-term spectrum of the matrix sentences. During an SRT measurement, the speech level^3^ was fixed at 50 dB SPL and the noise level varied adaptively using a one-down, one-up rule. The SRT was thus defined as the SNR at which listeners recognized 50% of full sentences (Levitt, 1971). Thirty sentences were presented to measure an SRT. The first 10 sentences were always the same (randomly presented) and were included to give listeners an opportunity to become familiar with the task. The initial SNR was 20 dB and it changed in 4-dB steps between sentences 1 and 14 and 2-dB steps between sentences 14 and 30. The SRT was calculated as the mean of the final 17 SNRs (the SNR for the 31st sentence was calculated and used in the SRT estimate but not actually presented). Each SRT estimate was measured three times and the mean was taken as the final SRT.

##### Conditions

SRTs were measured for three spatial configurations of target and masker sources: S_0_N_0_ with the speech and noise sources collocated in front of the listener at 0° azimuth; S_270_N_45_ with the speech and noise sources at 270° and 45° azimuth, respectively; and S_270_N_90_ with the speech and noise sources at 270° and 90° azimuth, respectively. These spatial configurations were selected as representative of conditions where the STOI and SNR evaluations predicted the benefit from the algorithm to be largest (S_270_N_45_), intermediate (S_270_N_90_), or none (S_0_N_0_) (see below).

In total, 36 SRTs were measured per participant: 2 listening modes (bilateral and unilateral) × 2 processing conditions (with and without processing) × 3 spatial configurations (S_0_N_0_, S_270_N_45_ and S_270_N_90_) × 3 SRT estimates per condition. Measurements were organized in three blocks of 12 SRT measurements each, one block per SRT estimate. Within each block, bilateral and unilateral listening modes were alternated and for each listening mode conditions were administered in random order. Participants did not know which condition they were being tested on.

##### Participants

Five NH adults (three women) participated in the tests. They were all native Spanish speakers. Their age range was 25 to 28 years, and their absolute hearing thresholds were less than 20 dB HL at the audiometric frequencies from 125 to 8000 Hz.

#### Sound localization tests

For NH listeners, sound source localization in a virtual horizontal plane was assessed for processed and unprocessed stimuli.

##### Stimuli

For each processing condition, localization was assessed for lowband (125-1000 Hz) and broadband (125-6000 Hz) noise bursts (four test conditions in total). Gaussian noise bursts were digitally generated and bandpass filtered (first-order Butterworth filter) to achieve the desired bandwidth. Stimuli had a duration of 200 ms (including 50-ms raised-cosine onset and offset ramps) and their level was fixed at 65 dB SPL. No level roving was applied to minimize the use of monaural level cues.

##### Procedure

The procedure was virtually identical to that of Lopez-Poveda et al. (2019). For each test condition, participants were presented with eight noise tokens for each one of the 13 azimuthal angles in the frontal horizontal plane from 270° to 90°, spaced every 15° (104 noise tokens in total). The 104 noise tokens were presented in random order. During the presentation of the stimuli, participants sat in front of a computer screen that displayed a top view of a human head with an array of speakers in front of the head. For each stimulus presentation, the subject was instructed to judge the azimuthal position of the sound source by clicking on the corresponding speaker on the computer screen. The click of a response triggered the processing of a freshly generated noise stimulus through the corresponding condition, and the presentation of the stimulus to the participant. The response screen displayed an array of 37 speakers spaced every 5° over an azimuth range from 270° to 90°, even though stimuli were presented every 15°. Feedback was not given to participants on the correctness of their responses.

Before data collection began, participants were encouraged to train themselves on the task by clicking on any of 37 speakers evenly spaced every 5° over an azimuth range from 270° to 90° and listening to the corresponding stimulus (that is, during training, participants could hear stimuli at all those azimuthal locations while for testing, stimuli were presented at a subset of locations). Training was provided independently for each test condition and for as long as each participant deemed necessary. The four test conditions were administered in random order and participants did not know which condition they were training on or being tested with.

##### Participants

Five NH adults (three women) participated in the experiment. Their age range was 26 to 31 years, and their absolute hearing thresholds were less than 20 dB HL at the audiometric frequencies from 125 to 8000 Hz. Three of these participants also participated in the speech-in-noise recognition test.

#### Equipment

The MATLAB software environment (R2017b, The Mathworks, Inc.) was used to perform all signal processing and implement all test procedures. All technical and experimental evaluations were conducted with digital stimuli sampled at a rate of 44100 Hz and with 24-bit resolution. For the experimental tests, stimuli were controlled using custom-made software and played via an RME Fireface UCX soundcard and presented to the listeners via ER-2 insert earphones (Etymotic Research Inc., Elk Grove Village, Illinois). Sound pressure levels (SPLs) were calibrated by placing the earphones in a Zwislocki DB-100 coupler connected to a sound level meter (Brüel&Kjaer, mod. 2238). Calibration was performed at 1 kHz and the obtained sensitivity was used at all other frequencies.

### RESULTS

#### SNR and STOI improvement for a single SSN source

**Figure 5** shows the SNR improvement (Eq. 11) provided by the algorithm in each ear, for a target speech source in competition with a single source of SSN, and for multiple azimuthal locations of the target and masker sources in the horizontal plane (0° elevation). The level of the target and the masker before HRTF filtering was −20 dB FS (i.e., 0 dB SNR).

**Figure 5.**
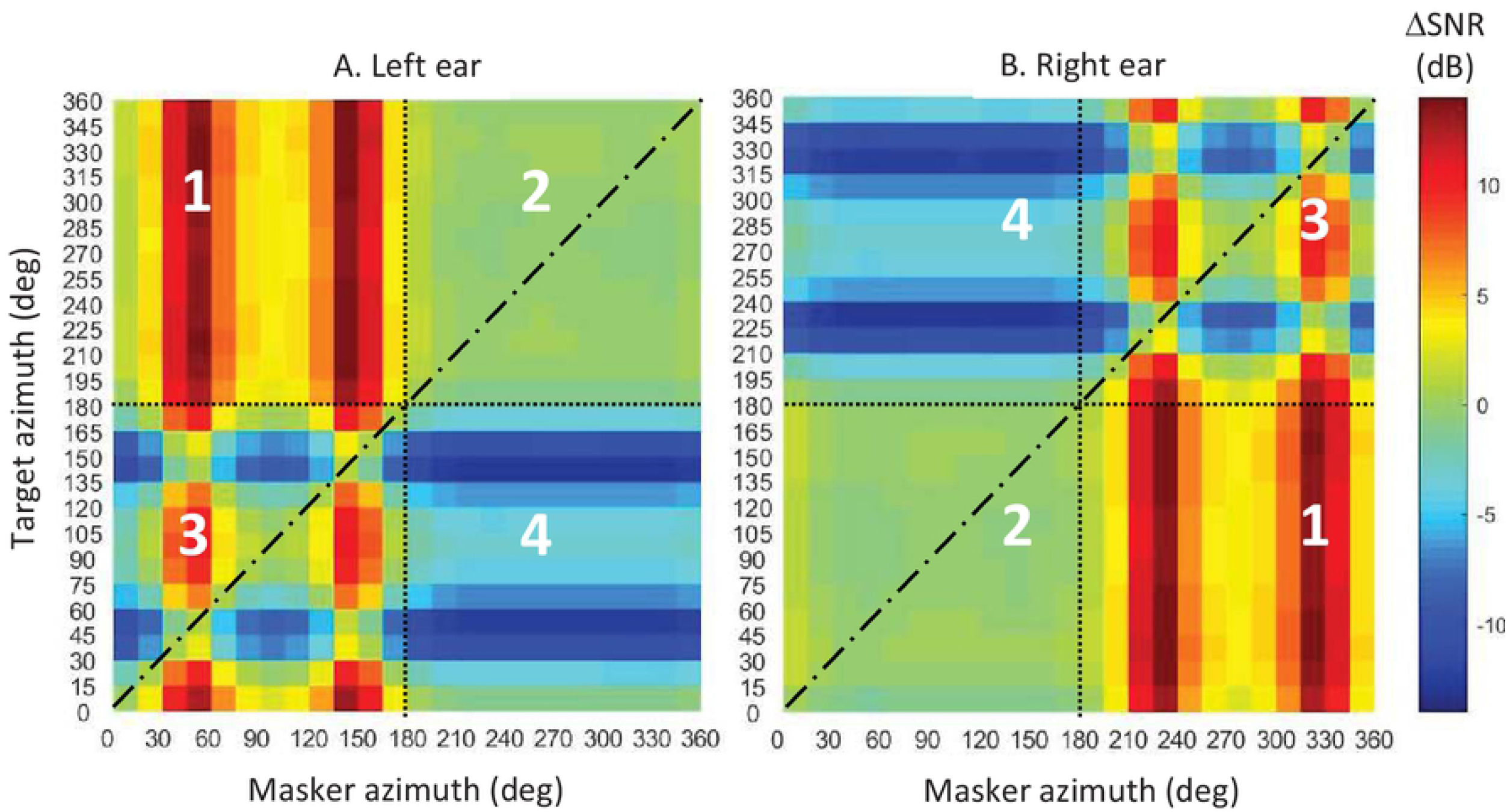
SNR improvement (ΔSNR) in the left and right ears provided by the algorithm with *β_diff_* and *α_diff_* for listening scenarios with the target and one masker source at azimuths from 0° to 360° every 15°. The color bar indicates the SNR improvement in decibels. Positive and negative values indicate higher and lower SNR for processed than for unprocessed stimuli, respectively. Dashed-dotted lines along the diagonals depict conditions with collocated target and masker sources. The dotted lines and numbers indicate the quadrants referred to in the main text.

The pattern of results in each panel (i.e., for each ear) suggests four quadrants, bounded by the dotted lines, and identified with numbers 1 to 4. Note that the quadrants are symmetrical for the left and right ears and so is the numbering. Quadrant 1 is for an ipsilateral target with a contralateral masker. In this case, the processor improves the SNR for most spatial configurations. The improvement can be as large as 14 dB and is caused by the attenuation of the contralateral masker (**Fig. 2**). Quadrant 2 is for an ipsilateral target and an ipsilateral masker. In this case, the processor hardly alters the SNR. Quadrant 3 is for a contralateral target with a contralateral masker. In this case, the processor can increase or decrease the SNR, depending on the actual azimuthal locations of the target and the masker. Quadrant 4 is for an ipsilateral masker with a contralateral target. In this case, the processor decreases the SNR overall because it attenuates the contralateral target (**Fig. 2**). Importantly, overall, the processor tends to improve the SNR in the ear ipsilateral to the target.

It can be mathematically demonstrated (not shown) that the algorithm does not alter the SNR in two special cases: (1) for collocated target and masker sources (i.e., along the dashed-dotted diagonals); and (2) for ipsilateral target and masker sources that are symmetrically placed with respect to the left-right axis (i.e., for a target at 45° and a masker at 135°).

**Figure 6** shows the STOI improvement (Eq. 12) provided by the algorithm with *β_diff_* and *α_diff_* for the same spatial locations of the target and masker sources as shown in **Fig. 5**. The pattern of results matches the pattern of SNR improvement illustrated in **Fig. 5**.

**Figure 6.**
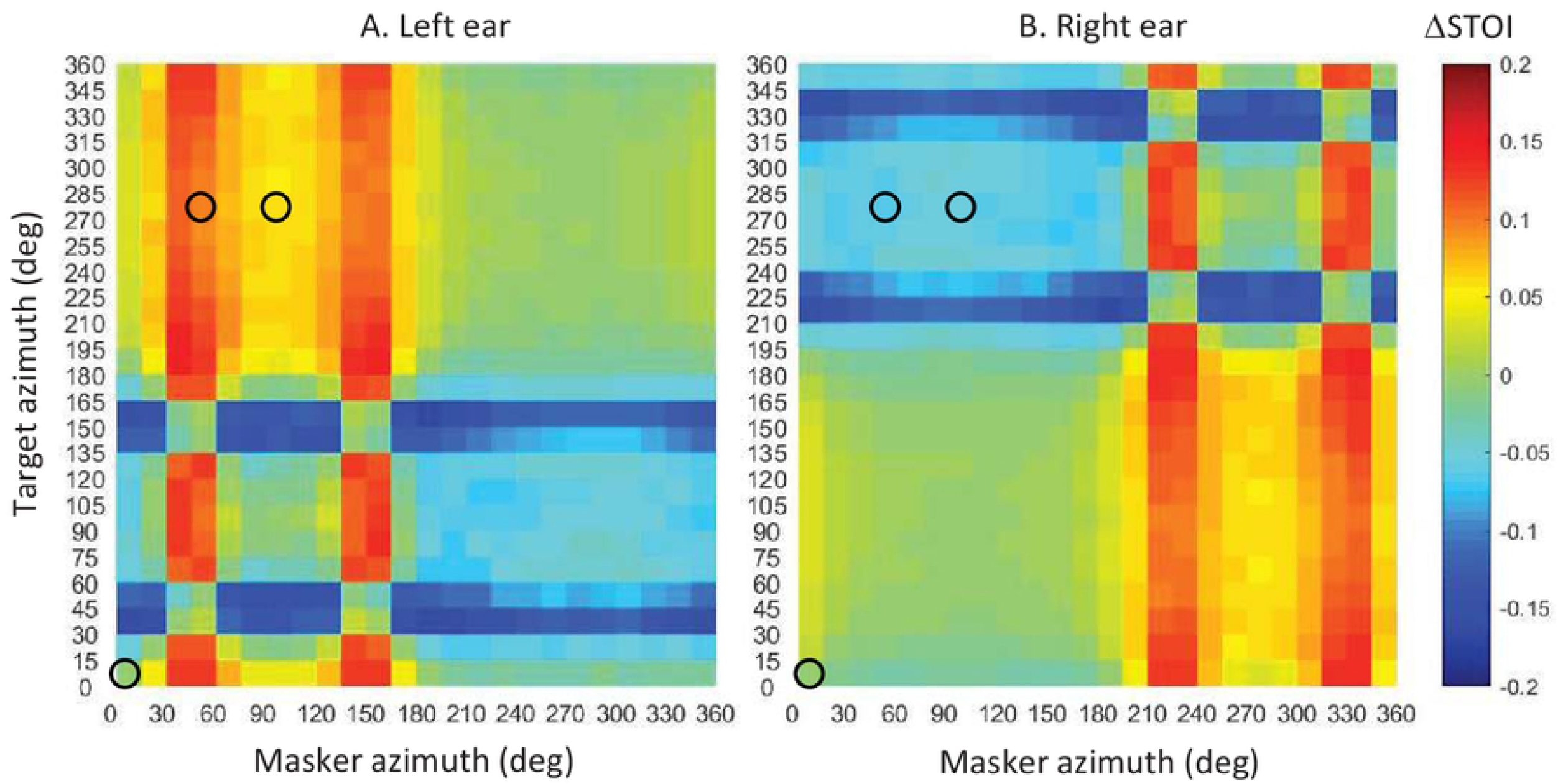
STOI improvement (ΔSTOI) for the left and right ears provided by the algorithm for various azimuthal locations of the target and masker sources in the horizontal plane. The SNR of the unprocessed stimuli before HRTF filtering was 0 dB. The color bar indicates the STOI improvement. Positive and negative values indicate higher and lower STOI for processed than for unprocessed stimuli, respectively. Circles depict the three target-masker spatial configurations (S_0_N_0_, S_270_N_45_, and S_270_N_90_) used in the experimental evaluation of the algorithm.

#### SNR and STOI improvement for multiple maskers and masker types

In the preceding section, it has been shown that the algorithm can improve the SNR and STOI in the ear ipsilateral to the target for simulated free-field listening conditions where the target speech source is in competition with a single source of SSN. **Figure 7** illustrates STOI and SNR improvements for listening conditions involving one or more simultaneous maskers and for three different masker types: IFFM, SSN, and WN. The improvement is shown only for the ear ipsilateral to the target, if any. For conditions involving multiple masker sources, the waveforms for all maskers were different. The only exception is indicated with the suffix “eq.” (for equal), where all masker sources had identical waveforms. For completeness, **Fig. 7** shows results for *α_diff_* and *β_diff_* calculated using individualized and non-individualized HRTFs (see Materials and Methods). In these calculations, the target and masker levels were set to −40 and −30 dB FS, respectively (i.e., the SNR was −10 dB).

**Figure 7.**
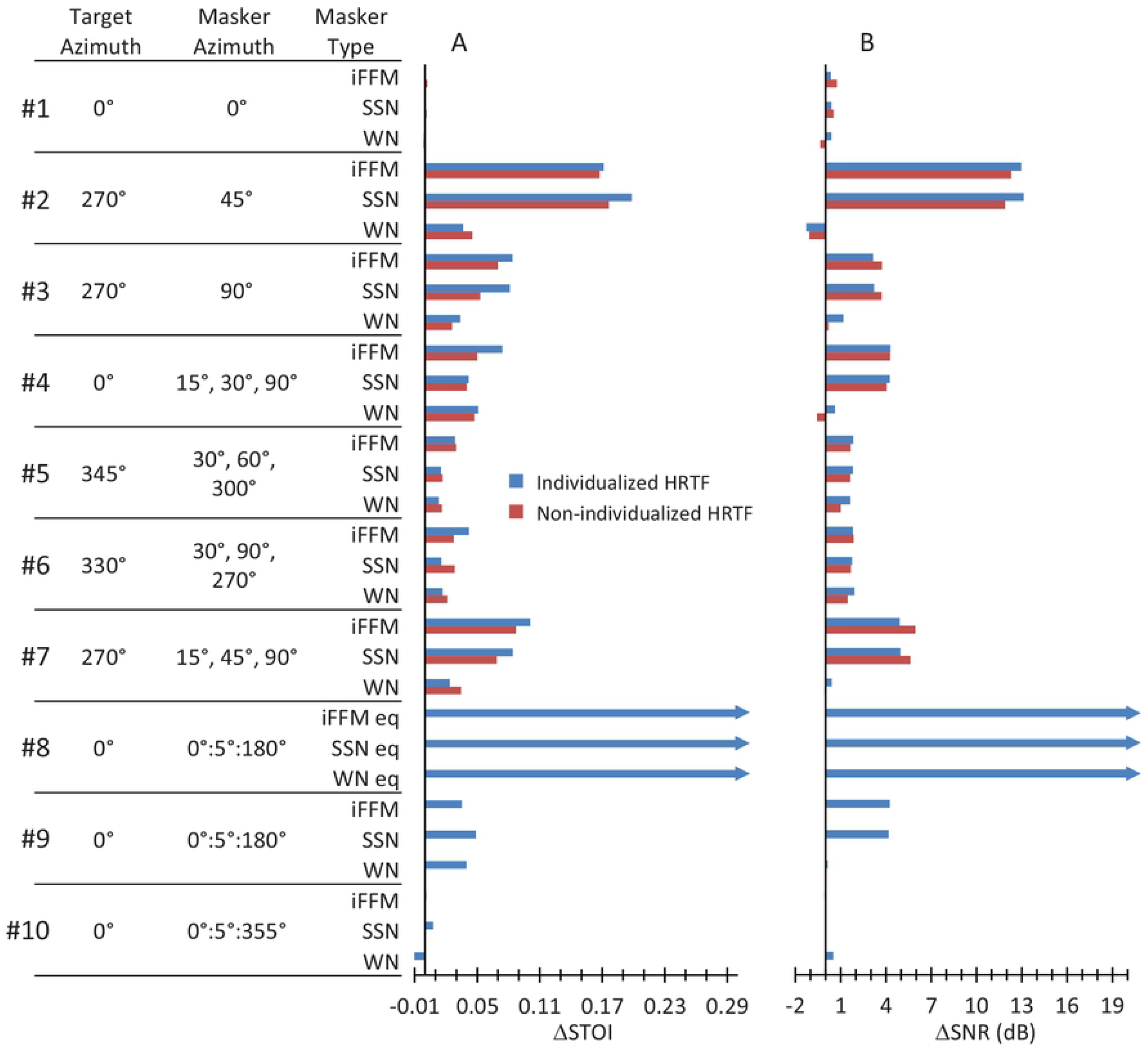
Improvement in STOI (**A**, ΔSTOI) and SNR (**B**, ΔSNR) in the left ear for 10 simulated listening scenarios with different number of maskers, and azimuth and masker locations. For each scenario, results are shown for three masker types (IFFM, SSN, WN). For scenarios #1 to #7, results are shown for *α_diff_* and *β_diff_* calculated using individualized and non-individualized HRTFs. Bars terminated with an arrow indicate that the actual bar extends beyond the corresponding x-axis range. Note that in some conditions, ΔSTOI and/or ΔSNR are equal to zero and hence the bar is not visible. See the main text for details.

The results in **Fig. 7** are grouped, from top to bottom, in 10 blocks of three masker types each. Blocks #1 to #3 involve a single masker, with target and masker azimuths as in the experimental evaluations (see below). Blocks #4 to #7 are for a single target source (at azimuths of 0°, 345°, 330°, and 270°) in competition with three simultaneous maskers at various locations, as indicated in the figure. Blocks #8 and #9 involve a target at 0° azimuth in competition with 37 identical (eq.) and different maskers, respectively, spanning the azimuth range from 0° to 180° (in 5° steps). Lastly, block #10 involves a target at 0° azimuth in competition with 72 different maskers spanning the azimuth range from 0° to 360° (in 5° steps).

**Figure 7** shows the following. (1) For most conditions, ΔSNR≥0 dB and ΔSTOI≥0 indicating that the algorithm improves the SNR and STOI. The exceptions were condition #1, with collocated target and masker sources, and condition #10, with maskers all around from 0° to 355°. (2) For every condition, the improvement is comparable for IFFM and SSN, and smaller for WN. (3) Improvements are overall larger when *α_diff_* and *β_diff_* are calculated using individualized than non-individualized HRTFs. However, important improvements are obtained even with non-individualized HRTFs. (4) The largest improvements occur for condition #8, with individualized HRTFs in a listening scenario with identical (equal) maskers on the contralateral hemifield, i.e., azimuths from 0° to 180°. This is not surprising because *α_diff_* and *β_diff_* were designed (Eq. 5) to produce perfect masker cancellation precisely in this hypothetical listening scenario.

#### Speech recognition in noise

While the SNR and STOI evaluations described earlier suggest that the proposed algorithm can improve the intelligibility of speech in competition with a single noise source, neither of them allows a direct assessment of the benefit provided by the algorithm in this task. For this reason, the algorithm was also evaluated experimentally by comparing SRTs for sentences in competition with a single SSN source. The evaluation was restricted to three spatial configurations of the target and masker sources (S_0_N_0_, S_270_N_45_, and S_270_N_90_), where the technical evaluations just described predicted the benefit to be none, large, and medium, respectively (see blocks #1 to #3 in **Fig. 7**).

**Figure 8A** allows a comparison of individual and group mean SRTs in bilateral listening for processed and unprocessed stimuli, for the three spatial configurations. Lower SRT values indicate better performance. **Figure 8B** shows the SRT improvement provided by the algorithm. A two-way RMANOVA was conducted to test for the effect of processing (processed, unprocessed) and spatial configuration (S_0_N_0_, S_270_N_45_, S_270_N_90_) on the SRT. The test revealed a significant main effect of processing [F(1,4)=218.26, p<0.001]. Mean SRTs (across locations and participants) were significantly better for processed than for unprocessed stimuli (−16.8 vs −13.6 dB SNR, respectively). It also revealed a significant main effect of spatial configuration [F(2,8)=1090.68, p<0.001]. Post-hoc pairwise tests with Bonferroni correction showed that SRTs were significantly worse (higher) in the collocated condition than in any of the spatially separated conditions [p(S_0_N_0_ vs S_270_N_45_)<0.001; p(S_0_N_0_ vs S_270_N_90_)<0.001], and they were significantly better in the S_270_N_45_ than in the S_270_N_90_ condition (p<0.001). The interaction between processor and spatial configuration was also significant [F(2,8)=138.31, p<0.001]. Post-hoc pairwise comparisons with Bonferroni correction showed that the algorithm improved SRTs significantly only in the S_270_N_45_ spatial configuration (p<0.001), but not in the S_0_N_0_ (p=0.290) or the S_270_N_90_ (p=0.063) configurations.

**Figure 8.**
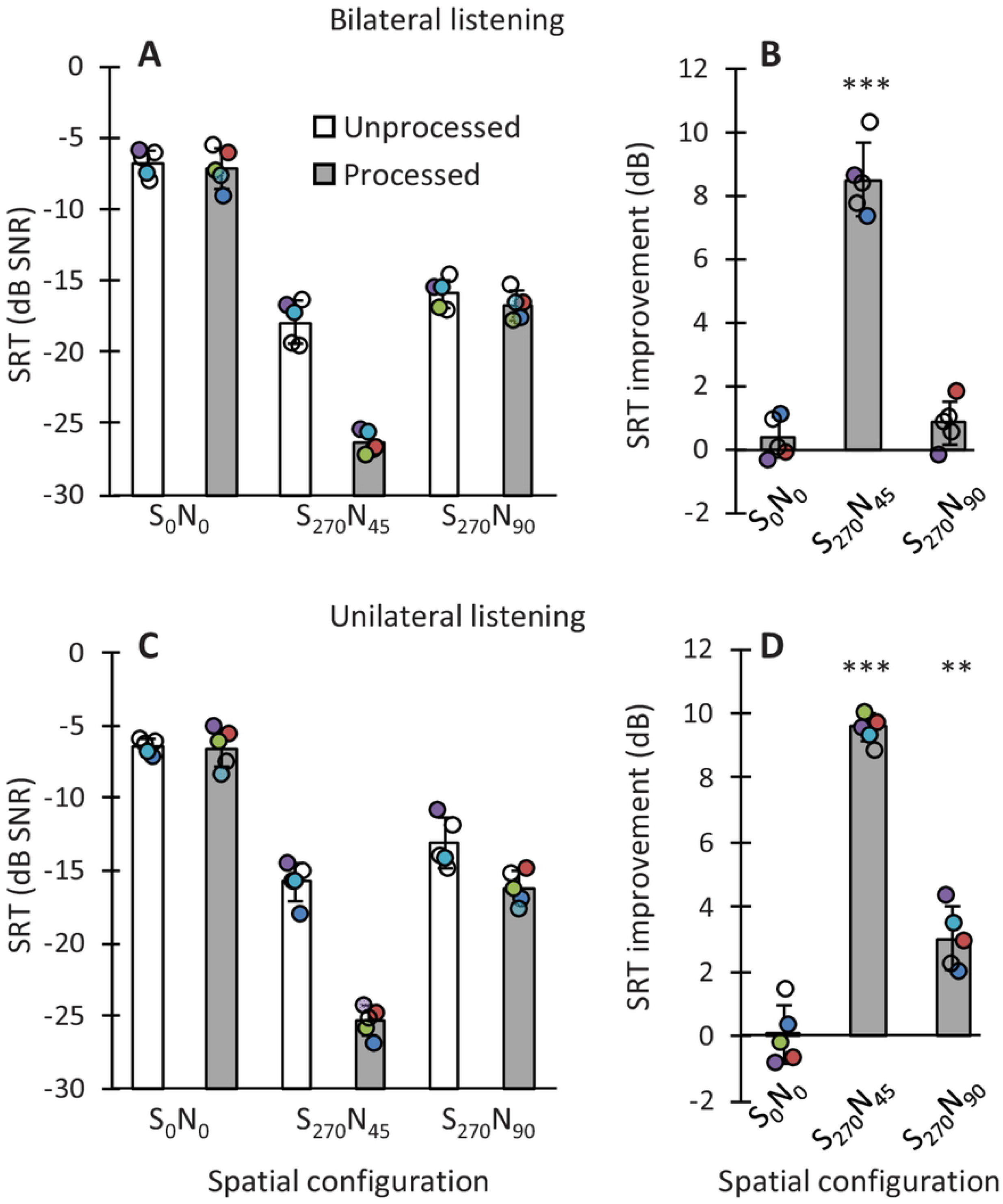
The effect of processing on speech-in-noise recognition for selected target-masker spatial configurations (S_0_N_0_, S_270_N_45_, and S_270_N_90_), in unilateral and bilateral listening modes. **A, C.** Speech reception thresholds for processed (grey bars) and unprocessed stimuli (white bars) in bilateral and unilateral listening, respectively. **B, D.** SRT improvement provided by the processor. In all panels, circles illustrate individual data, bars illustrate group mean scores (N=5), and error bars illustrate plus and minus one standard deviation. Unilateral listening involved listening with the left ear (i.e., the ear ipsilateral to the target source in the S_270_N_45_, and S_270_N_90_ configurations). ** *p*≤0.01, *** *p*≤0.001.

**Figure 8C-D** show the effect of the algorithm on SRTs when listening with the left ear alone, i.e., the processing was binaural, but stimuli were presented to the ear ipsilateral to the target source. A two-way RMANOVA revealed a significant main effect of processing [F(1,4)=577.4, p<0.001]. Mean SRTs (across locations and participants) were significantly better for processed than for unprocessed stimuli (−16.0 vs −11.8 dB SNR, respectively). It also revealed a significant main effect of spatial configuration [F(2,8)=801.63, p<0.001]. Post-hoc pairwise tests with Bonferroni correction showed that SRTs were significantly worse (higher) in the collocated condition than in any of the spatially separated conditions [p(S_0_N_0_ vs S_270_N_45_)<0.001; p(S_0_N_0_ vs S_270_N_90_)<0.001], and they were significantly better in the S_270_N_45_ than in the S_270_N_90_ condition (p<0.001). The interaction between processor and spatial configuration was also significant [F(2,8)=158.99, p<0.001]. Post-hoc pairwise comparisons with Bonferroni correction showed that the algorithm improved SRTs significantly in the S_270_N_45_ (p<0.001) and S_270_N_90_ (p=0.002) but not in the S_0_N_0_ (p=0.886) conditions.

#### Sound source localization

As suggested by the directivity patterns of **Fig. 2**, the algorithm with *β_diff_* and *α_diff_* can attenuate contralateral low frequencies (<1500 Hz) and slightly enhance high frequencies (>1500 Hz), distorting head-shadow ILDs. **Figure 9** illustrates this effect more clearly by illustrating the level at each ear and the corresponding ILD as a function of sound source azimuth for white (0-22050 Hz), broadband (125-6000 Hz), and lowband (125-1000 Hz) noises of −20 dB FS. Note that the algorithm distorts the stimulus level, particularly in the contralateral ear, and that the distortion is comparatively greater for lowband noise than for broadband or white noises.

**Figure 9.**
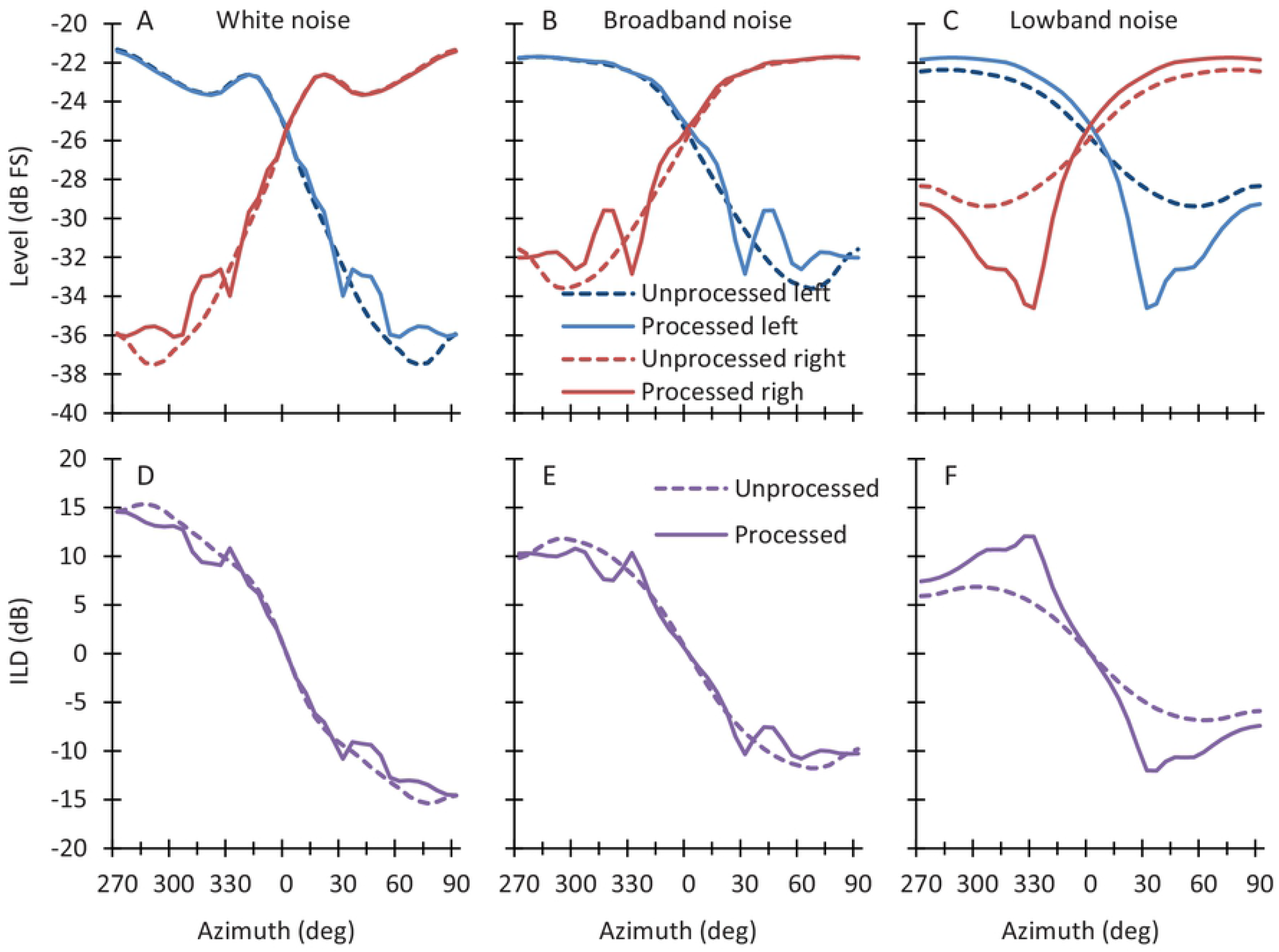
**Top.** The level at the left and right ears for unprocessed and processed stimuli. The stimulus level before HRTF filtering was −20 dB FS. **Bottom.** Inter-aural level difference (ILD) for unprocessed and processed stimuli. Each column is for a different noise bandwidth, as indicated at the top: white (0-22050 Hz), broadband (125-6000 Hz), and lowband (125-1000 Hz).

To assess the potential impact of these distortions, sound source location was assessed for unprocessed and processed lowband (LBN) and broadband (BBN) noises. Performance was quantified using the root mean square angle error between presentation and response angles. **Figure 10** shows a comparison of individual and group mean angle error for the four conditions. Smaller values indicate better performance. A two-way RMANOVA was conducted to test for the effects of the processing (processed and unprocessed) and noise type (LBN and BBN) on localization. The RMANOVA revealed that neither the processing [F(1,4)=0.50, p=0.52] nor the noise bandwidth [F(1,4)=0.37, p=0.58] had a statistically significant main effect on the angle error. The RMANOVA, however, revealed a statistically significant interaction between processing and noise type on angle error [F(1,4)=38.8, p=0.003]. Post-hoc pairwise comparisons with Bonferroni correction, however, revealed that the algorithm did not have a significant effect on the angle error for LBN (p=0.720) or BBN (p=0.100).

**Figure 10.**
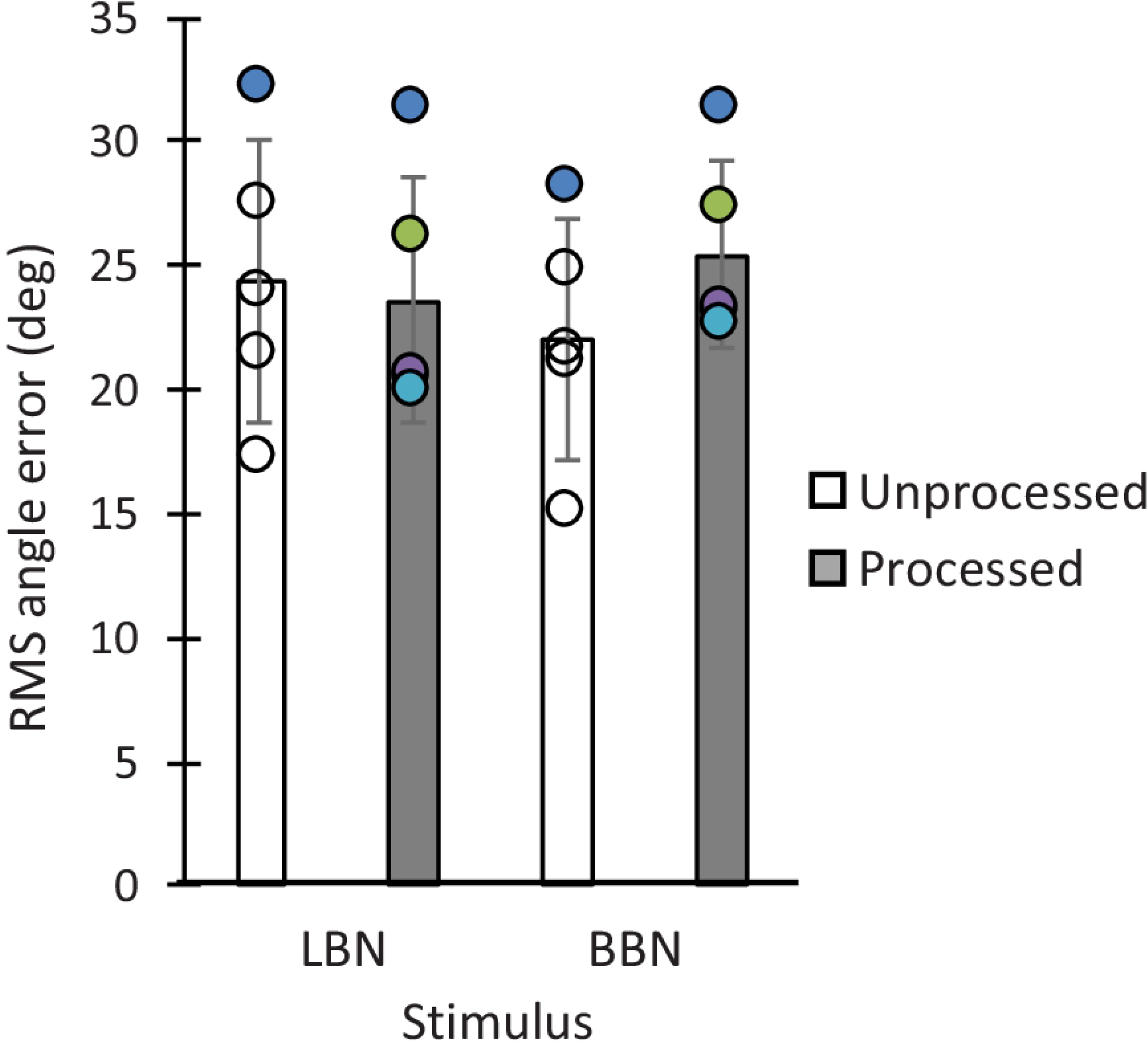
The effect of processing on the localization of lowband (LBN) and broadband (BBN) noise bursts in a virtual horizontal plane. Open and filled bars illustrate group mean angle error scores for unprocessed and processed stimuli, respectively. Error bars illustrate plus and minus one standard deviation of the group mean. Circles illustrate individual scores.

### DISCUSSION

We have proposed a pre-processing algorithm (Eq. 3) to attenuate the contralateral sound field in each ear by weighted subtraction of the contralateral stimulus. The weights for the left and right ear [*β*(*f*) and *α*(*f*)] are regarded as free parameters. Here, we have set them equal to *β_diff_* and *α_diff_*, respectively, and calculated them with Eq. (5) using KEMAR HRTFs. With these weights, the algorithm can improve the SNR (**Figs. 5 and 7**) and the intelligibility of speech in competition with one (**Figs. 6 and 8**) or more interfering sound sources (**Fig. 7**), with minimal impact on sound source localization (**Fig. 10**).

#### The choice of weights *α*(*f*) and *β*(*f*)

The specific weights employed here, *β_diff_* and *α_diff_*, were chosen ad hoc with the aim to cancel a ‘diffuse’ masker in the contralateral hemifield, spanning the azimuthal range from 0° to 180° (see Eq. 5). These weights, however, need not be optimal or suitable for all applications. Indeed, the algorithm produces different SNR improvements depending on the weights. **Figure 11** shows that for a target at 0° azimuth in competition with a single SSN source at azimuths from 0° to 360°, the SNR improvement in the left ear would be different depending on the value of *β*(*f*). In these examples, the values of *β*(*f*) were calculated using Eq. (5b) but averaging HRTFs over different azimuth ranges, as indicated by the inset (note that *β_diff_* involved averaging KEMAR HRTFs over an azimuth range from 0° to 180°). The figure shows that while *β_diff_* causes SNR improvements to peak at masker azimuths of 45° and 135°, other *β*(*f*) values can produce larger improvements and at different masker azimuths.

**Figure 11.**
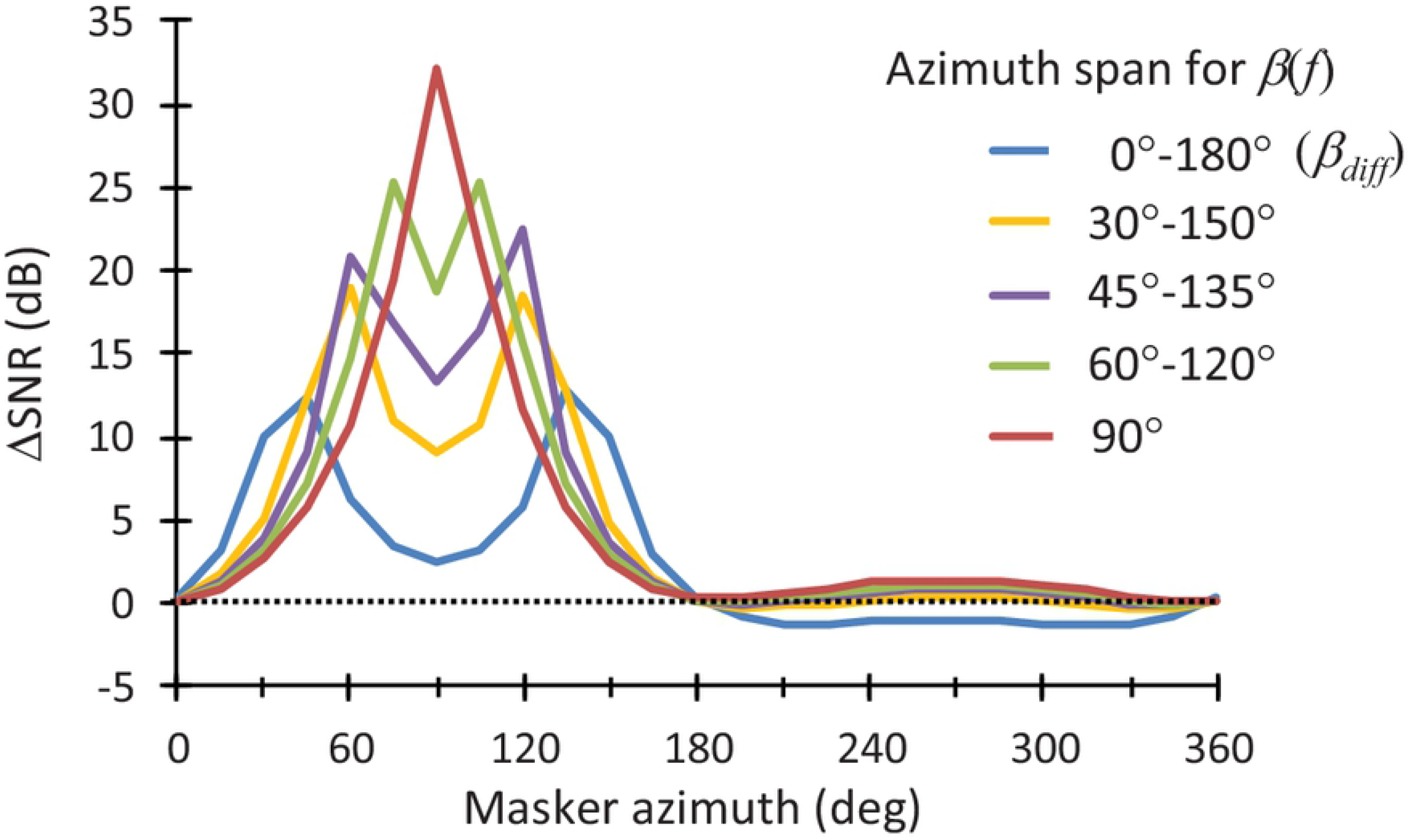
SNR improvement produced by the algorithm in left ear for a target at 0° azimuth in competition with a SSN source at azimuths from 0° to 360° (in steps of 15°). Each trace illustrates the improvement for different values of *β*(*f*) calculated using Eq. (5a) but averaging KEMAR HRTFs over different azimuth ranges, as indicated by the inset. The improvement is relative to HRTF-filtered but otherwise unprocessed stimuli. See the main text for details.

On the other hand, *α_diff_* and *β_diff_* were appropriate to produce the sought effect of attenuating contralateral maskers but only when maskers were low frequencies. With *α_diff_* and *β_diff_*, however, the algorithm enhanced contralateral high-frequency maskers slightly (**Fig. 2**). As a result, *α_diff_* and *β_diff_* produce larger SNR improvements for maskers with relatively weaker than stronger high-frequency content (> 1500 Hz). Because white noise has more high-frequency content than IFFM or SSN, this explains why ΔSNR and ΔSTOI tended to be smaller for white noise than for IFFM or SSN (see **Fig. 7** and also compare **Fig. 3** vs. **Fig. 4**). Setting *α_diff_* and *β_diff_* equal to zero for frequencies above 1500 Hz would prevent the amplification of contralateral high-frequencies, which would extend the benefit from the algorithm to maskers with strong high-frequency content, including white noise.

#### Relation with binaural unmasking

The present algorithm was conceived as a pre-processing tool to improve the SNR. However, we incidentally found that it can coarsely simulate binaural intelligibility differences (BILDs). The BILD is defined as the SRT improvement that results from having dichotic rather than diotic speech-in-noise stimuli (Levitt and Rabiner, 1967). In free-field listening, it may be defined as the SRT improvement (decrease) for any target-masker spatial configuration relative to the S_0_N_0_ condition, where stimuli are almost diotic (note that diotic stimuli is unlikely to occur in natural listening situations due to small asymmetries between the ears). Because BILDs exceed the benefit provided by the head shadow alone, i.e., the benefit from listening with the ear that has the better acoustic SNR, (e.g. Bronkhorst and Plomp, 1998), the current view is that BILDs also reflect some form of binaural interaction in the central auditory system to ‘cancel’ the masker(s) (e.g., Durlach, 1963; Culling, 2007; Hauth and Brand, 2018).

**Figure 12A** shows experimental BILDs for a target at 0° azimuth in competition with a single SSN source as a function of masker azimuth (denoted as the S_0_N_X_, with X indicating the masker azimuth). **Figures 12B and 12C** show BILDs for configurations that included a second SSN source at 105° azimuth (S_0_N_105_N_X_), and a third source at 255° azimuth (S_0_N_105_N_255_N_X_), respectively. All data are from Peissig and Kollmeier (1997). For comparison, **Figure 12D-F** show the SNR improvements relative to the S_0_N_0_ condition produced by the present algorithm for corresponding listening conditions and for two sets of weights: *α_diff_* and *β_diff_*, and *α*_270±30_ and *β*_90±30_. The former were calculated using Eq. 5 by averaging KEMAR HRTFs over the azimuth range 0°-180° for *β*(*f*) and 180°-360° for *α*(*f*), while the latter were calculated using Eq. (5) but averaging KEMAR HRTFs over a narrower azimuth range of 60°-120° for *β*(*f*) and 240°-300° for *α*(*f*). Also shown are SNR improvements for unprocessed stimuli, i.e., improvements produced by the head-shadow alone. Plots show the SNR in the ear with the largest SNR for any given masker azimuth, which is equivalent to assuming that listeners use the ear with the better acoustic SNR to recognize the speech. As the figure shows, the algorithm with weights *α*_270±30_ and *β*_90±30_ produces SNR improvements that are larger but roughly similar to the pattern of experimental BILDs. In addition, SNR improvements are much larger for processed than for unprocessed stimuli.

**Figure 12.**
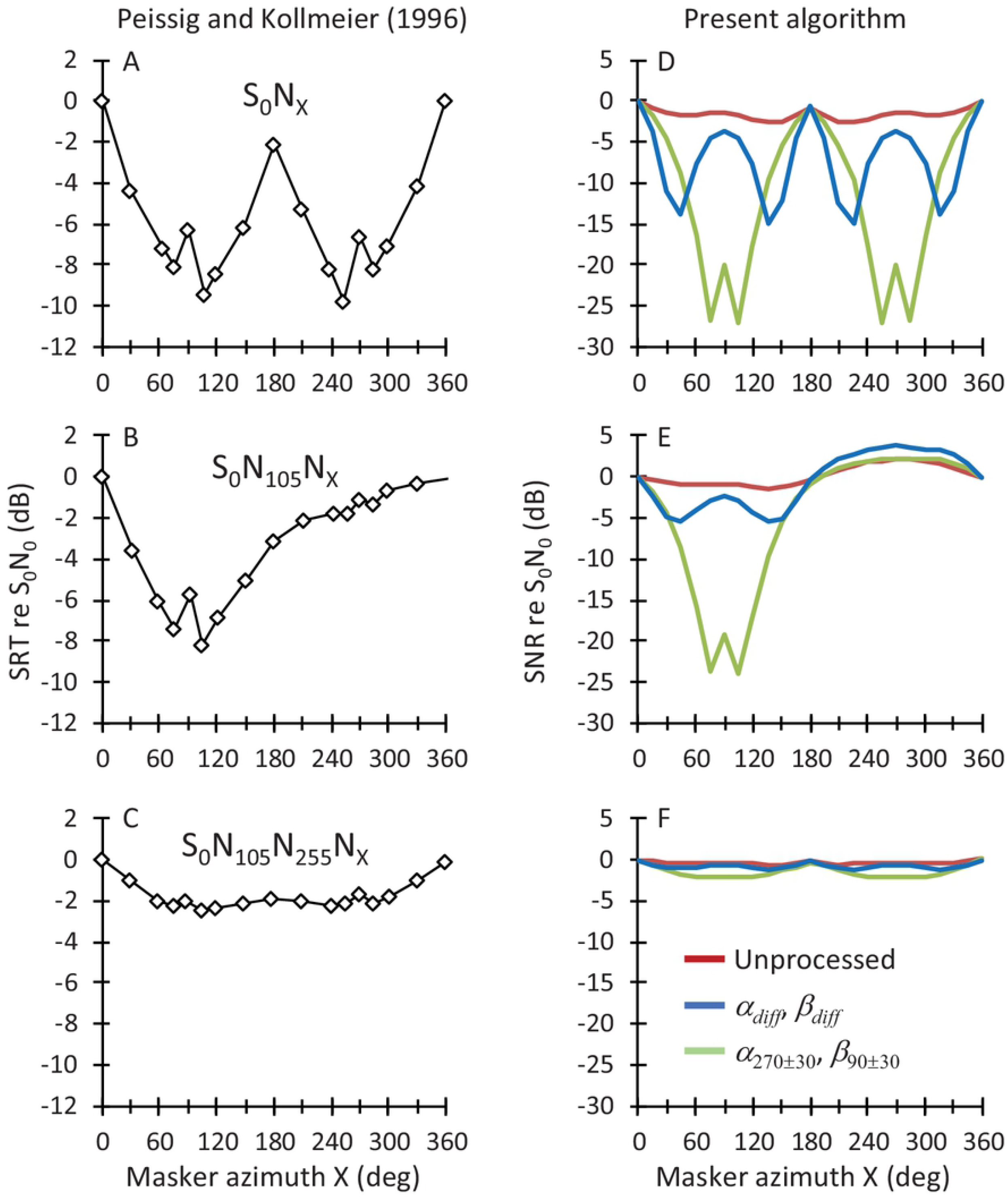
Comparison of the SNR provided by the present algorithm in the ear with the largest SNR (**right panels**) with experimental SRTs (data from Peissig and Kollmeier, 1997) (**left panels**) relative to a spatial configuration with the target and masker collocated at 0° azimuth. Lower values indicate better performance. Note the different ordinate scales in the left and right panels. The top, middle, and bottom rows show results for a target at 0° azimuth in competition with one, two, and three different SSN sources, respectively. The abscissa illustrates the azimuth location (X) of one of the SSN sources. The middle and bottom panels illustrate the results when the second SSN source, and third SSN source were located at 105° and 255° azimuth, respectively. See the main text for details.

Based on the similarity between the pattern of SNR improvements and experimental BILDs in **Fig. 12**, it is tempting to speculate that the central auditory system could weight and subtract the stimuli at the two ears, and that the proposed algorithm could inspire binaural intelligibility models. The idea of binaural subtraction is not new and goes back to the equalization-cancellation theory of Durlach (1963). He theorized that “the auditory system attempts to eliminate the masking component by first transforming the stimuli presented to the two ears so as to equalize the two masking components, and the subtracting”. A main difference with Durlach’s theory is that the present algorithm involves fixed weighting of the stimuli presented to the two ears rather than equalization of the masking components at the two ears. Furthermore, the present findings suggest that the weighting might be based on the interaural ratio of HRTFs averaged over an appropriate azimuth range. Further research is necessary to investigate these ideas, and explore the potential of the present algorithm as a model of binaural intelligiblity.

#### Limitations

The measured SRT improvements were generally smaller than predicted by ΔSNR. For example, the measured mean SRT improvement in the S_270_N_45_ condition was ~8.5 dB, thus smaller than one would expect from **Fig. 5A** (~14 dB) if listeners used the ear with the better acoustic SNR to perform the SRT task. Likewise, the measured SRT improvement in the S_270_N_90_ was also smaller than predicted by ΔSNR in **Fig. 5A** (0.8 vs ~4.0 dB). The reason is uncertain. Perhaps, listeners did not use the ear with the better SNR to perform the speech recognition task. Or, perhaps, the processed stimuli interfered binaurally in the auditory brain of the listeners to prevent them from taking full advantage of the SNR provided by the algorithm in the ear ipsilateral to the target (e.g., Jerger et al., 1993; Mussoi and Bentler, 2017). The latter explanation is partially supported by the fact that the measured SRT improvements were greater, and closer to ΔSNR values, in unilateral than in bilateral listening (compare **Fig. 8B** with **Fig. 8D**). The latter explanation is, however, insufficient because the observed SRT improvements in unilateral listening were still smaller than predicted by ΔSNR. Whatever the reason, the present findings show that it would be wrong to expect actual SRT improvements to be as large as ΔSNR.

As explained in the Introduction, pre-processing algorithms like the one proposed here are often intended to improve the SNR to the users of assistive listening devices. The present tests were limited to listeners with normal-hearing listeners. Further research is required to investigate the benefits provided by the present algorithm when used in combination with hearing aids and/or cochlear implants.

#### Final remarks

Given that listeners can use better-ear glimpsing in cocktail party scenarios (Brungart and Iyer, 2012), and that the proposed algorithm can improve the SNR in the ear ipsilateral to the target source, the algorithm can improve the intelligibility of speech in competition with one or more sound sources without the user’s intervention, without making any assumption about the temporal or spectral characteristics of the target and masker sources, and without making assumptions about the specific location of the interfering sound sources. In broad terms, it functions as a binaural beamformer that attenuates (or cancels) the contralateral sound field (**Fig. 2**), and whose frequency-dependent directivity pattern depends on the weights *α*(*f*) and *β*(*f*) (**Fig. 11**). It operates in the frequency domain and its implementation in a hearing device would require two microphones (one per ear), and a means of exchanging the stimulus spectrum across the ears. Overall, the present algorithm is conceptually simpler and arguably easier to implement than many SNR enhancement pre-processing approaches (reviewed by Hamacher et al., 2005; Baumgärtel et al. 2015a; see the Introduction), thus potentially useful for implementation in binaural hearing devices.

Dieudonné and Francart (2018) proposed a single-delay-and-subtract algorithm “to effectively enhance low-frequency ILDs without the need of estimations of auditory cues or distorting the incoming sound” (their p. 79). Their algorithm effectively operates as a binaural beamformer that attenuates the contralateral sound field (see their Fig. 1B). On the other hand, the equalization-cancellation model was conceived to explain binaural masking level differences. In contrast with those approaches, the present algorithm was conceived as a practical solution to the general problem of cancelling out as many maskers as possible in cocktail party listening scenarios without having to know *a priori* the spectra or the locations of the target or the maskers (see section entitled The Algorithm). Given the disparity in motivations, it is remarkable that the present algorithm operates as a binaural beamformer that enhances head-shadow ILDs, as the algorithm of Dieudonne and Francart (2018) does, and that when set with appropriate values for *α* and *β* (**Fig. 12**), it qualitatively simulates binaural unmasking in the free field, as the equalization-cancellation model does.

### CONCLUSIONS

1. The proposed algorithm (binaural weighted subtraction of the contralateral stimulus) attenuates contralateral sounds, which in free-field listening conditions improves the SNR in the ear ipsilateral to the target sound source.
2. The subtraction weights may be set as needed. However, setting them equal to the ratio of ipsilateral to contralateral HRTFs averaged over an appropriate azimuth range can improve the SNR by as much as 20 dB, and in multiple listening scenarios.
3. Binaural weighted subtraction can improve speech reception thresholds in noise by up 8.5 dB in bilateral listening and 10 dB in unilateral listening.
4. Binaural weighted subtraction using HRTF-based weights qualitatively accounts for binaural unmasking when speech is presented in competition with multiple maskers and for multiple target-masker spatial arrangements.
5. Binaural HRTF-weighted subtraction is simple and effective, which makes it potentially suitable for implementation in binaural hearing devices.

## Authors contributions

EALP designed the algorithm and the study. AEM performed the technical evaluations and wrote the software tools for the experimental evaluations. FMSV collected the experimental data with human participants. EALP wrote the manuscript. All authors analyzed the data and revised the manuscript.

## ACKNOLWEDGEMENTS

We thank Milagros J. Fumero for help with data collection, and Miriam I. Marrufo-Pérez, Peter T. Johannesen and Thibaud Leclere for useful discussions. Work supported by the Spanish Ministry of Science and Innovation (grant PID2019-108985GB-I00) and by MED-EL GmbH (Innsbruck, Austria).

## APPENDIX. THE ALGORITHM IN DETAIL

In a listening scenario where the target source with spectrum, *S*(*f*), is presented in competition with multiple (*n*) maskers with spectra *M*_1_(*f*), *M*_2_(*f*),…, *M_n_*(*f*), the stimulus spectrum in the left and right ears, *L*(*f*) and *R*(*f*), can be calculated as follows:

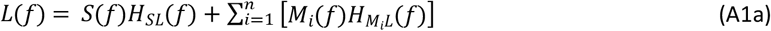

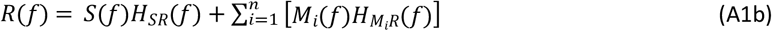

where *f* denotes frequency (Hz), and *H_SL_*(*f*), *H_M_i_L_*(*f*) are the complex HRTFs for the target (*S*) and the *i*-th masker (*M_i_*) for the left (*L*) ear, and *H_SR_*(*f*) and *H_M_i_R_*(*f*) are the corresponding HRTFs for the right (*R*) ear.

Substituting Eqs. (A1a) and (A1b) into Eq. (1a) and Eq. (1b), we get:

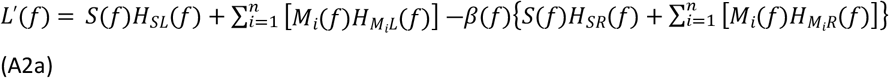

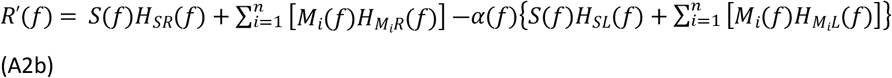

For convenience, Eqs. (A2a) and (A2b) may be re-written as follows:

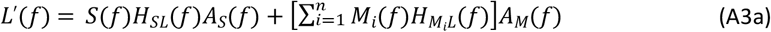

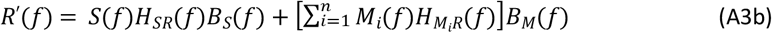

Where:

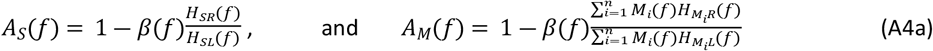

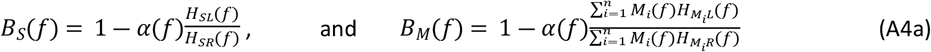

The right-hand side of Eqs. (A3a) and (A3b) contain one term for the target, *S*(*f*), plus one term for the maskers, *M_i_*(*f*). Perfect cancellation of the masker(s) at the two ears can be achieved by making the masker terms equal to zero. That is, perfect cancellation of the masker(s) in the left ear can be achieved by choosing *β*(*f*) such that *A_M_*(*f*) = 0 for all frequencies. Similarly, perfect cancellation of the maskers in the right ear can be achieved by choosing *α*(*f*) such that *B_M_*(*f*) = 0 for all frequencies.

From Eq. (A4a), it can be easily shown that *A_M_*(*f*) = 0 if and only if:

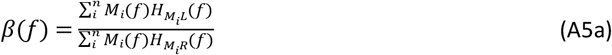

Similarly, from Eq. (A4b), *B_M_*(*f*) = 0 if and only if:

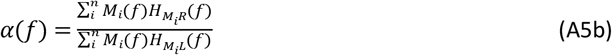

Note that:

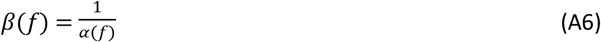

## Footnotes

1 For sound sources in the left hemifield, the algorithm improves the SNR in the left ear by attenuating sound sources in the right hemifield. For sources in the right hemifield, the algorithm improves the SNR in the right by attenuating sound sources in the left hemifield. The listener can then attend to the rightear or the left-ear stimulus, which ever has the higher SNR for the target, depending on whether the target is in the right or the left hemifield.

2 http://recherche.ircam.fr/equipes/salles/listen/download.html

3 The speech level was lower than used in most conventional tests (50 rather than 65 dB SPL) because the technical evaluation of the algorithm predicted that SRTs in noise would improve up to 14 dB SNR in some conditions, and hence the noise level at the SRT would have been high (~90 dB SPL) and uncomfortable if the speech level had been 65 dB SPL.

